# FKBPL and FKBP8 regulate DLK degradation and neuronal responses to axon injury

**DOI:** 10.1101/2021.08.20.457064

**Authors:** Bohm Lee, Yeonsoo Oh, Eunhye Cho, Aaron DiAntonio, Valeria Cavalli, Jung Eun Shin, Yongcheol Cho

## Abstract

DLK is a key regulator of axon regeneration and degeneration in response to neuronal injury. To understand the molecular mechanisms controlling the DLK function, we performed yeast two-hybrid screening analysis and identified FKBPL as a DLK-binding protein that bound to the kinase domain and inhibited the kinase enzymatic activity of DLK. FKBPL regulated DLK stability through ubiquitin-dependent DLK degradation. We tested other members in the FKBP protein family and found that FKBP8 also induced DLK degradation as FKBPL did. We found that Lysine 271 residue in the kinase domain of DLK was a major site of ubiquitination and SUMO3-conjugation and responsible for FKBP8-mediated degradation. In vivo overexpression of FKBP8 delayed progression of axon degeneration and neuronal death following axotomy in sciatic and optic nerves, respectively, although axon regeneration efficiency was not enhanced. This research identified FKBPL and FKBP8 as new DLK-interacting proteins that regulated DLK stability by MG-132 or bafilomycin A1-sensitive protein degradation.

## Introduction

Dual leucine zipper kinase (DLK), an upstream MAP triple kinase, is a key regulator of neuronal damage responses playing a major role in axon regeneration and degeneration (Holland *et al*, 2016b; Welsbie *et al*, 2018). DLK regulates the JNK signaling pathway when neurons are under stress conditions and is essential for injury-induced retrograde signaling that induces injury-responsive differential gene expression (Shin *et al*, 2012a, 2019). DLK is also required for axon degeneration because genetic depletion of DLK impairs axon degeneration (Miller *et al*, 2009). Therefore, finding molecular regulators of DLK and understanding their regulatory mechanisms are required for better understanding axon regeneration and degeneration.

Protein degradation pathways are a core axis regulating neuronal responses to a diverse range of stresses (Nakata *et al*, 2005; Collins *et al*, 2006). Since DLK is a key regulator of neurodegenerative signal transduction, the post-translational modification of DLK has been studied intensively to understand the mechanisms balancing DLK activity, localization, and protein levels (Huntwork-Rodriguez *et al*, 2013; Watkins *et al*, 2013; Larhammar *et al*, 2017; Holland *et al*, 2016). Modifications such as phosphorylation and palmitoylation are essential for the regulation of DLK functions and stability across different cellular contexts (Montersino & Thomas, 2015; Niu *et al*, 2020; Martin *et al*, 2019). DLK protein levels are modulated by stress signaling through the PHR1 E3 ubiquitin ligase and the de-ubiquitinating enzyme USP9X, which is a key pathway determining neuronal fates after injury (Babetto *et al*, 2013; Larhammar *et al*, 2017; Watkins *et al*, 2013; Huntwork-Rodriguez *et al*, 2013). Therefore, identifying the molecular mechanism controlling DLK stability with its interaction of DLK-regulating proteins is required to understand the DLK-mediated signal pathway.

In the present study, we report that FKBPL was a new DLK-binding protein identified by yeast two-hybrid screening. FKBPL is a member of the FK506-binding protein (FKBP) family of immunophilins, the group of the conserved proteins binding with immunosuppressive drugs, such as FK506, rapamycin and cyclosporin A. FKBPL bound to the kinase domain of DLK and inhibited its kinase activity. FKBPL induced DLK degradation by ubiquitin-dependent pathway. Comparative analysis showed that another FKBP member, FKBP8, also bound to DLK and induced DLK degradation as FKBPL did. We found that the lysine residue at position 271 in the kinase domain was a target site for DLK ubiquitination that was responsible for MG-132 and bafilomycin A1-sensitive DLK degradation. The lysine residue also served as SUMO3-conjugating site, implying that the evolutionarily conserved lysine residue in the kinase domain was a potential site for regulating DLK functions. FKBP8 promoted lysosomal degradation of ubiquitinated DLK, while a K271R mutant form of DLK was resistant to FKBP8-mediated DLK degradation. In vivo overexpression of FKBP8 delayed axon degeneration in sciatic nerves after axotomy and exhibited a protective effect in retinal ganglion cells after optic nerve crush.

Because DLK mediates retrograde signals to the nucleus to determine neuronal fates for regeneration or apoptosis after axonal injury (Shin *et al*, 2012a; Watkins *et al*, 2013), identifying proteins regulating DLK protein levels are important for understanding mechanisms of DLK-regulated axon regeneration or DLK-induced neuronal death. Here, we present new DLK-interacting molecules, FKBPL and FKBP8, that regulate DLK degradation by ubiquitin-dependent lysosomal pathway.

## Results

### FKBPL was identified as a DLK-binding protein

To identify DLK-interacting proteins, we performed yeast two-hybrid screening analysis and found *Fkbpl*, *Tedc2*, and *Tuft1* as potential candidates. By analyzing two independent reporter systems, we confirmed the *Fkbpl* gene product as the protein that is most stably bound to DLK (Figures 1A and 1B). FKBPL is a member of FK506-binding proteins (FKBPs), a family of proteins that bind to tacrolimus (FK506), an immunosuppressant molecule, and have prolyl isomerase activity (Siekierka *et al*, 1989). The co-immunoprecipitation assay validated the screening result and showed that exogenously expressed DLK and FKBPL in HEK293T cells were stably associated (Figure 1C).

**Figure 1.**
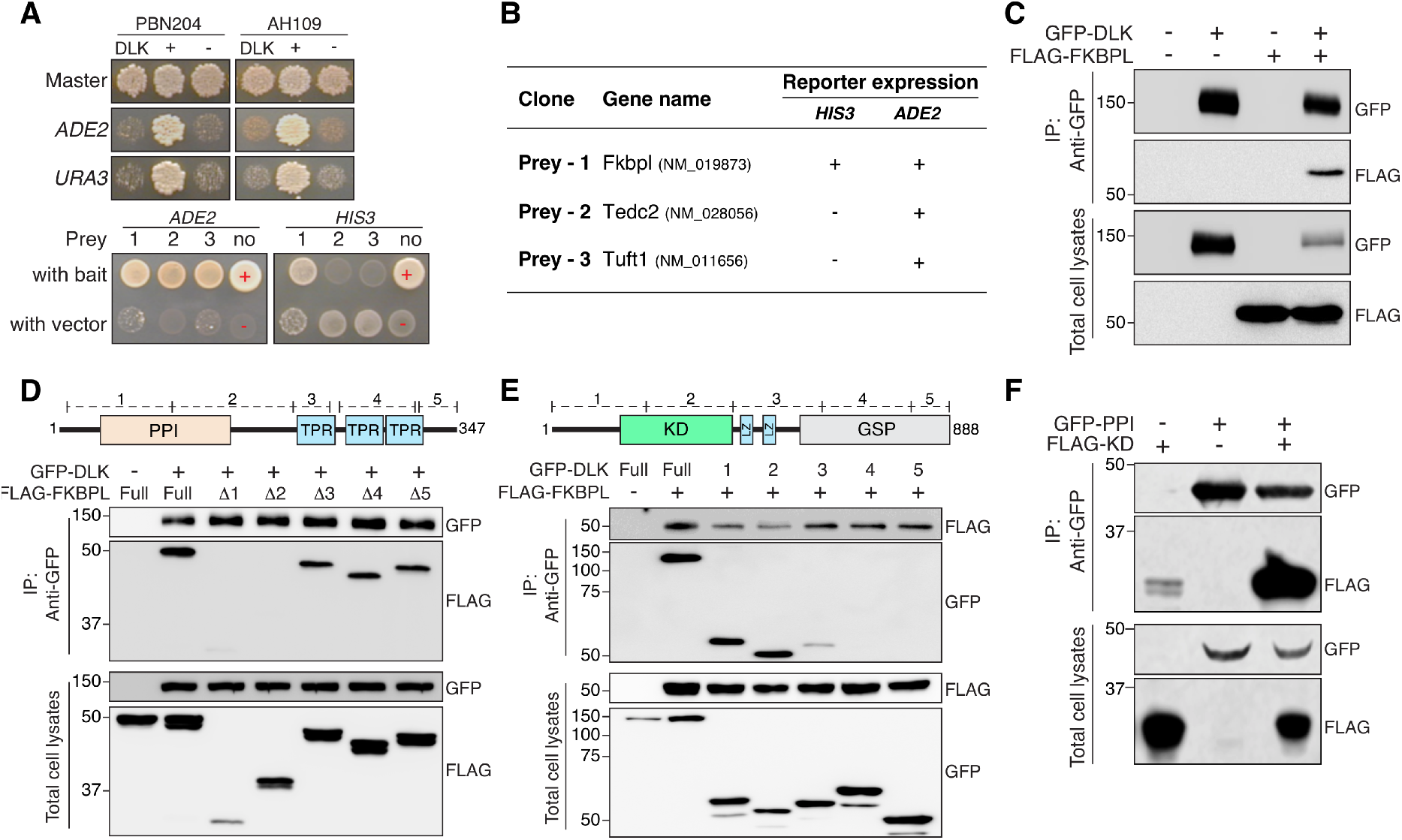
The PPI domain of FKBPL and the kinase domain of DLK are responsible for their interaction. (A) Yeast two-hybrid screening analysis was performed to identify DLK-interacting proteins in two different yeast strains PBN204 and AH109. Master plates indicated the positive controls for the selection. *ADE2* and *URA3* indicated -Ade and -Ura minimal medium. It validated that DLK itself did not activate *ADE2* nor *URA3* genes (Top). The ADE2-selection screening identified three preys (1, *Fkbpl*; 2, *Tedc2*; 3, *Tuft1*). ADE2-minimal medium selection with an additional selection of HID3-showed that prey 1 survived and made the colony. Red + indicated the positive control identical with the top panel. (B) The *Fkbpl* gene product exhibits the most stable association with DLK of the three potential candidates by two reporter-analysis systems. (C) Western blot analysis of the co-immunoprecipitation of DLK and FKBPL in HEK293T cells. GFP-DLK and FLAG-FKBPL were co-transfected to HEK293T cells. The protein lysates were immunoprecipitated using an anti-GFP antibody followed by SDS-PAGE analysis. (D) Schematic diagram of the protein domains of FKBPL and western blot analysis for the co-immunoprecipitation analysis of DLK with FKBPL-deletion mutants (PPI; peptidylprolyl isomerase domain, TPR; tetratricopeptide repeat domain). Anti-GFP antibody was used for immunoprecipitation. (E) Schematic diagram of the protein domains of DLK and western blot analysis for the co-immunoprecipitation analysis of FKBPL with partial DLK proteins (KD; kinase domain, LZ; leucine zipper motif, GSP; Gly, Ser, and Pro-rich domain). Anti-GFP antibody was used for immunoprecipitation. (F) Western blot analysis for the co-immunoprecipitation of the PPI domain and KD. Anti-GFP antibody was used for immunoprecipitation.

To map the region responsible for the interaction, FKBPL-deletion mutants were subjected to co-immunoprecipitation analysis. The deletion of the N-terminal part of FKBPL, including the peptidylprolyl isomerase (PPI) domain, impaired the interaction between DLK and FKBPL (Figure 1D). In addition, the N-terminal region of DLK, including the kinase domain, mediated the interaction because a partial form of DLK with its N-terminal region intact was still able to bind to FKBPL (Figure 1E). These results indicated that the N-terminal regions of both proteins were responsible for their interaction. Because the PPI and the kinase domains formed a stable association, they served as the interface for their interaction (Figure 2F). The yeast two-hybrid screening and the co-immunoprecipitation analyses revealed that FKBPL was a new binding partner of DLK and their interaction was mediated by the PPI domain of FKBPL and the kinase domain of DLK.

**Figure 2.**
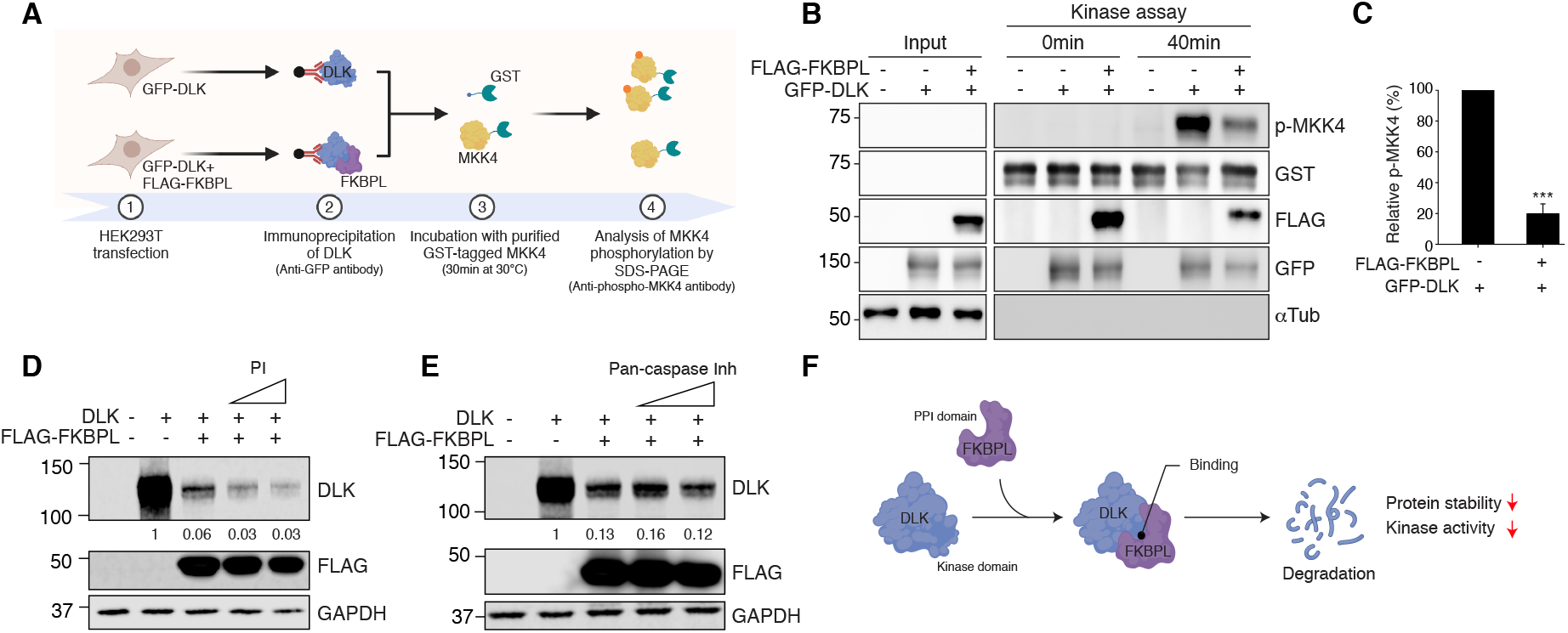
FKBPL inhibited the kinase activity of DLK and lowered the DLK protein level. (A) Illustration of DLK in vitro kinase assay. GFP-DLK was expressed from HEK293T cells with or without FKBPL co-expression. Immunopurified GFP-DLK was incubated in the reaction buffer including the substrate of purified GST-MKK4. (B) Western blot analysis for in vitro kinase assays. Phosphorylated MKK4 level was visualized by anti-phospho-MKK4 antibody. Anti-GST, FLAG, GFP antibody was used for detecting GST-MKK4, FLAG-FKBPL, and GFP-DLK protein from input and kinase reaction samples. Anti-α-tubulin antibody was used for the internal control. (C) Quantification of the relative p-MKK4 from (A) (n=3 for each condition; \*\*\**p*<0.001 by t-test; mean ± S.E.M.). (D and E) Western blot analysis for GFP-DLK and FLAG-FKBPL protein level with protease inhibitor (PI) (D) or pan-caspase inhibitor (Pan-caspase Inh) (E) treatment at different doses. (F) Proposed model of the interaction between DLK and FKBPL.

### FKBPL inhibited the kinase activity of DLK and induced degradation

Since FKBPL interacted with DLK via its kinase domain, we performed in vitro kinase assay to investigate whether DLK kinase activity was modulated by its association with FKBPL (Holland *et al*, 2016) (Figure 2A). Incubating the DLK substrate GST-MKK4 with immunopurified GFP-DLK induced phosphorylation of GST-MKK4 in vitro (Figure 2B). However, MKK4 phosphorylation was significantly reduced when GST-MKK4 was incubated with immunopurified GFP-DLK associated with FLAG-FKBPL (Figures 2B and 2C), indicating that the association of DLK and FKBPL inhibited the DLK kinase activity. As FKBPL bound to the kinase domain of DLK, the association of FKBPL might physically block the interaction of its substrate to the kinase domain.

DLK kinase activity was required for its stabilization and introducing mutations impairing the kinase activity caused destabilization of DLK at the protein level (Huntwork-Rodriguez *et al*, 2013). Since FKBPL bound to DLK and inhibited its kinase activity, we tested the protein level of DLK with FKBPL co-expression and found that the DLK protein level was significantly lowered when FKBPL was co-expressed in HEK293T cells (Figure 2D). However, the FKBPL-mediated DLK reduction was not prevented by applying an inhibitor of enzymatic activity of a broad spectrum of serine proteases, cysteine proteases, metalloproteases, and calpains (Figure 2D). In addition, an inhibitor of caspases had no effect on preventing the DLK reduction (Figure 2E). Therefore, the direct cleavages from proteases might not be responsible for the FKBPL-induced DLK degradation. These results showed that FKBPL was a negative regulator of DLK and inhibited DLK kinase activity.

### FKBPL and FKBP8 induced lysosomal DLK degradation

FKBPs are a group of proteins containing an FK506-binding domain and a peptidylprolyl isomerase (PPI) domain (Ghartey-Kwansah *et al*, 2018; Heitman *et al*, 1992; Kang *et al*, 2008; Schmid *et al*, 1993). Since FKBP family shares similar structures with high sequence homology (Tong & Jiang, 2015), we expanded the analysis to other FKBP proteins to test the reduction of DLK protein levels. Considering the DLK functions regulating axon regeneration and degeneration in dorsal root ganglion neurons, we reviewed the abundance of FKBPs’ mRNA in mouse L4,5 DRG tissues and mouse sciatic nerves from our previously pulished datasets (Shin *et al*, 2019; Shin *et al*, 2018a; Lee *et al*, 2021). To consider the neuronal expression of the FKBPs, the microarray dataset from cultured mouse embryonic DRG neurons was presented as the relative sizes of the circles (Figure 3A). The analysis showed that the mRNA levels of FKBP4, FKBP8 and FKBP12 were relatively higher than the others from adult mouse DRGs and sciatic nerves (Figure 3A). In addition, the neuronal expression profiles of FKBP mRNAs showed that FKBP4, −8 and −12 were relatively higher than the others, which was analyzed from the cultured embryonic DRG neurons without non-neuronal cells (Cho *et al*, 2013). This analysis implied that FKBP4, −8 and 12 were the potential candidates of the DLK-regulating FKBPs in sensory neurons.

**Figure 3.**
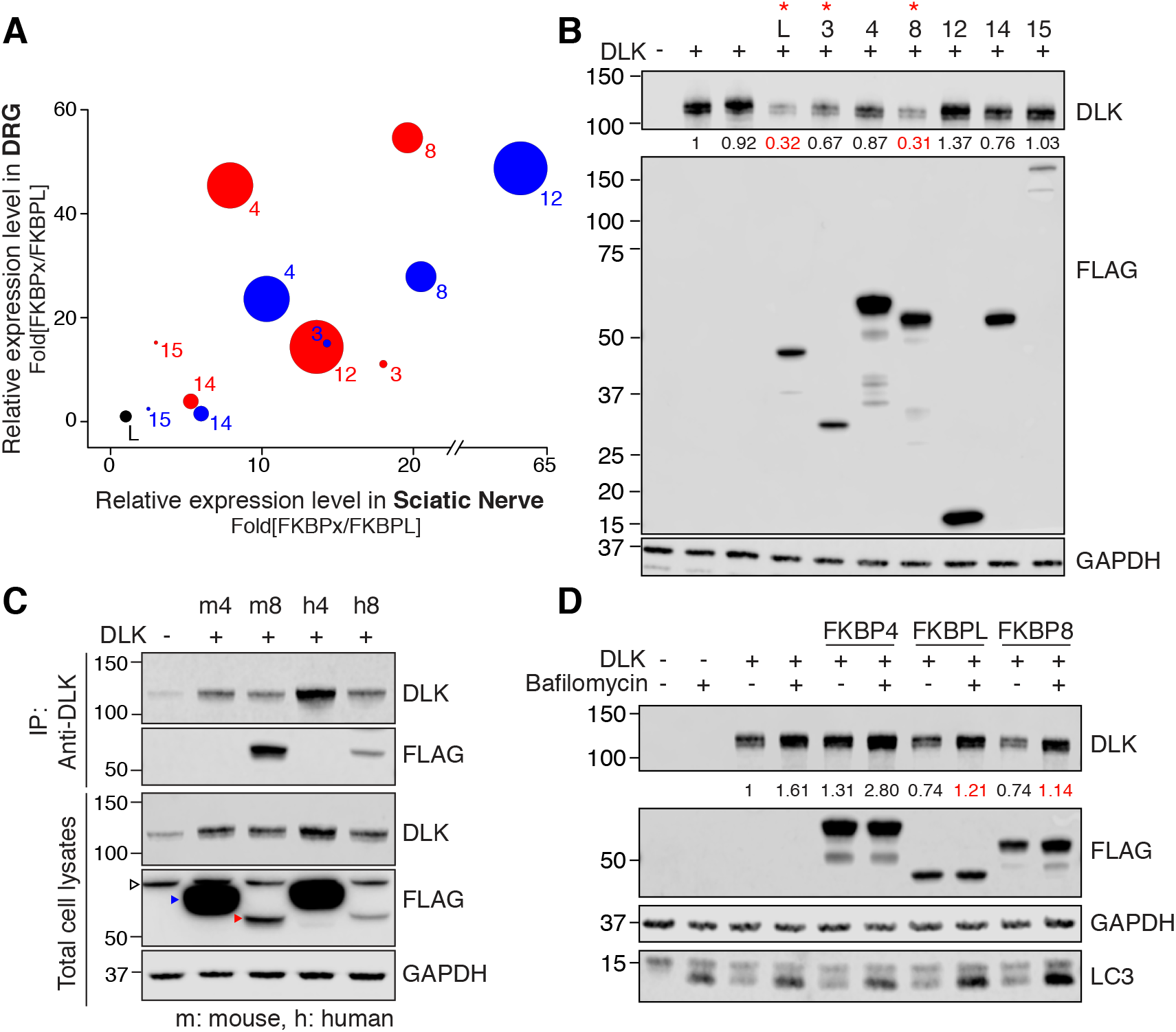
FKBPL and FKBP8 induced lysosome-dependent DLK degradation. (A) Comparative analysis of relative expression levels in mouse DRG (Shin *et al*, 2019; Lee *et al*, 2021), sciatic nerve tissue (Shin *et al*, 2018b), and cultured embryonic DRG neurons (Cho *et al*, 2013). Red and blue circles indicated Illumina short-read sequencing and Nanopore direct RNA long-read sequencing, respectively. Circle sizes indicated relative levels of micro array data from cultured embryonic DRG neurons. (B) Western blot analysis for the expression of DLK with FKBPs. The numbers indicate normalized relative intensity. (C) Western blot analysis for the immunoprecipitation of DLK with a mouse (m) and human (h) FKBP4/8. Empty arrowhead, non-specific band; blue arrowhead, FKBP4; red arrowhead, FKBP8. (D) Western blot analysis for the expression of DLK and FKBPL/4/8 with or without bafilomycin A1 treatment. The numbers indicate the normalized relative intensity.

Next, we tested DLK destabilization with co-expression of the FKBPs. FKBPL co-expression in HEK293T cells lowered DLK protein level to 32% (Figure 3B). The western blot analysis showed that FKBP8 was the most potent for lowering DLK protein levels similar to FKBPL, whereas FKBP4 had no significant effect on DLK protein reduction (Figures 3B and 3D). In addition, co-immunoprecipitation result showed that both human and mouse FKBP8 was associated with DLK, while FKBP4 did not interact with DLK (Figure 3C).

FKBP8 is known to regulate Parkin-independent mitophagy, which involves the removal of mitochondria via autophagy and lysosomal degradation (Yoo *et al*, 2020; Misaka *et al*, 2018; Lim & Lim, 2017; Bhujabal *et al*, 2017; Kang *et al*, 2008). As FKBP8 bound to DLK and lowered DLK protein levels, we tested if FKBP8- or FKBPL-induced DLK reduction was mediated by lysosomal degradation. When HEK293T cells were incubated with bafilomycin A1, FKBPL- and FKBP8-induced DLK degradation was inhibited indicating that FKBPL- and FKBP8-induced DLK degradation was regulated by lysosomal protein degradation functions. Moreover, the basal DLK protein level without co-expressing FKBP8 or FKBPL was increased by bafilomycin A1 treatement, suggesting that the baseline DLK turnover is mediated by lysosomal degradation (Figure 3D). These results showed that DLK protein levels were regulated by lysosomal functions and FKBPL- and FKBP8-induced DLK protein degradation.

### The kinase domain was a major target of DLK ubiquitination and sumoylation

DLK protein degradation is regulated by PHR1 E3 ligase (Collins *et al*, 2006; Nakata *et al*, 2005). As FKBPL- or FKBP8-induced DLK protein reduction required lysosomal degradation function, we tested if FKBP8-dependent DLK protein reduction was regulated by the ubiquitin-dependent protein degradation pathway. First, DLK was subjected to ubiquitination assay, which showed that HA-epitope tagged ubiquitin proteins were covalently conjugated with DLK protein in HEK293T cells (Figure 4A). When FKBP8 was co-expressed with DLK, ubiquitinated DLK proteins were significantly reduced, which was reversed by incubating transfected HEK293T cells with MG-132, an inhibitor of ubiquitin-dependent protein degradation (Figure 4A). This data indicated that FKBP8-induced DLK degradation via ubiquitin-dependent protein degradation pathway and FKBP8 overexpression accelerated degradation of ubiquitinated DLK protein.

**Figure 4.**
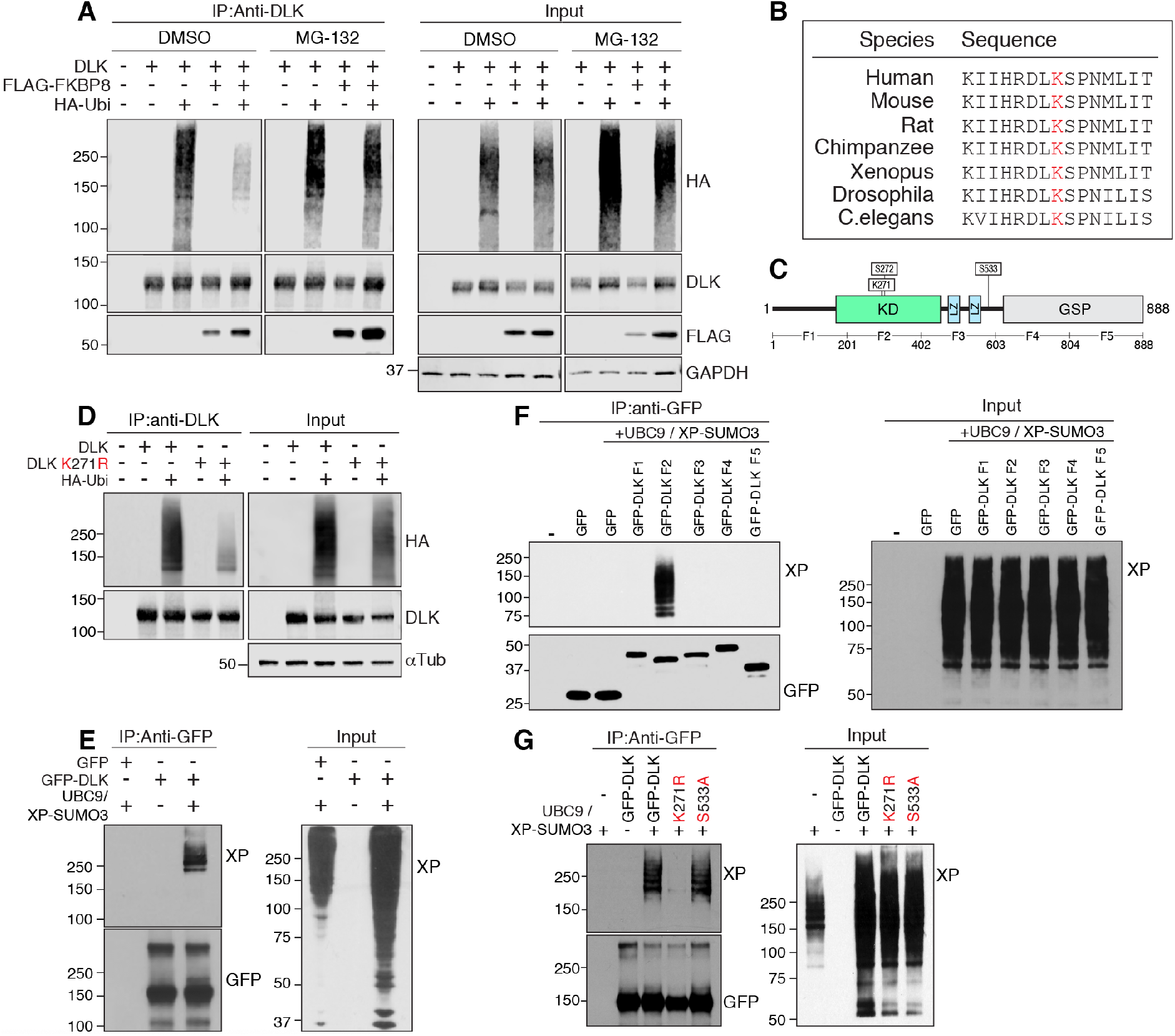
Lysine 271 in the kinase domain is a major target of DLK ubiquitination and sumoylation. (A) Western blot analysis for immunoprecipitation assays of DLK and HA-ubiquitin (HA-Ubi) with or without MG-132 treatment. (B) Alignment of amino acid sequences adjacent to the lysine 271 in the kinase domain of DLK in various species (Red; lysine at 271). (C) Schematic diagram of DLK protein domains (KD; kinase domain, LZ; leucine zipper motif, GSP; Gly, Ser, and Pro-rich domain). (D) Western blot analysis for immunoprecipitation assays with DLK, a K271R mutant, and HA-ubiquitin (HA-Ubi). (E) Western blot analysis for SUMO-denaturation immunoprecipitation assays with GFP-DLK, UBC9, and XP-SUMO3. (F) Western blot analysis for SUMO-denaturation immunoprecipitation assay with partial DLK proteins. (G) Western blot analysis for SUMO-denaturation immunoprecipitation assays with DLK, a K271R mutant, and an S533A mutant.

To identify lysine residues responsible for ubiquitination, we searched the lysine residues in the kinase domain of DLK because FKBPL interacted with the DLK kinase domain. The amino acid sequences alignment showed that DLK kinase domain sequences were evolutionarily conserved with high homology (Figure 4B). From the aligned lysine residues, we recognized that K271 was followed by S272, the serine residue known to be phosphorylated by JNK and critical for the DLK kinase activity because DLK^S272A^ mutant had no kinase activity (Huntwork-Rodriguez *et al*, 2013). In addition, S272 residue was one of the top three sites for JNK-dependent phosphorylation of DLK and the only serine residue in the kinase domain among them (Figure 4C). Moreover, this region was highly conserved across species (Figure 4B). Therefore, we hypothesized that this region controls DLK protein function, for example, via regulating its kinase activity and protein stability, and that K271 served as a potential lysine residue for post-translational modifications, including ubiquitination.

To test the residue, DLK^K271R^ mutant was subjected to ubiquitination assay, which showed that substitution of lysine at 271 to arginine significantly reduced the efficiency of ubiquitin conjugation to DLK^K271R^ mutant (Figure 4D). This indicated that the lysine at 271 was a major target site of DLK ubiquitination. Furthermore, this residue was responsible for DLK SUMOylation because DLK was a SUMO3-conjugating target protein and only the partial form of DLK including the kinase domain could be fully SUMOylated (Figures 4E and 4F). However, introducing mutation at K271 to arginine dramatically impaired SUMO3 conjugation (Figure 4G). These results revealed that the K271 site was the major ubiquitination target site and the SUMO3-conjugating lysine site.

### FKBP8 mediated ubiquitin-dependent DLK degradation

Because K271 was responsible for DLK ubiquitination, we investigated whether this site was required for FKBP8-induced DLK degradation. The western blot analysis showed that the DLK^K271R^ mutant was resistant to FKBP8-induced degradation (Figure 5A). Introducing a substitution mutation at K271 inhibited FKBP8-dependent DLK degradation (Figure 5B). This result indicated that ubiquitnated DLK at K271 was the major target of the FKBP8-mediated degradation pathway and suggested that ubiquitinated DLK protein was recruited to FKBP8-mediated protein degradation complex. Notably, DLK^K271R^ displayed a less efficient association with FKBP8, implying that lowering the DLK ubiquitination efficiency resulted in less interaction with FKBP8 protein and its associated complex (Figure 5C).

**Figure 5.**
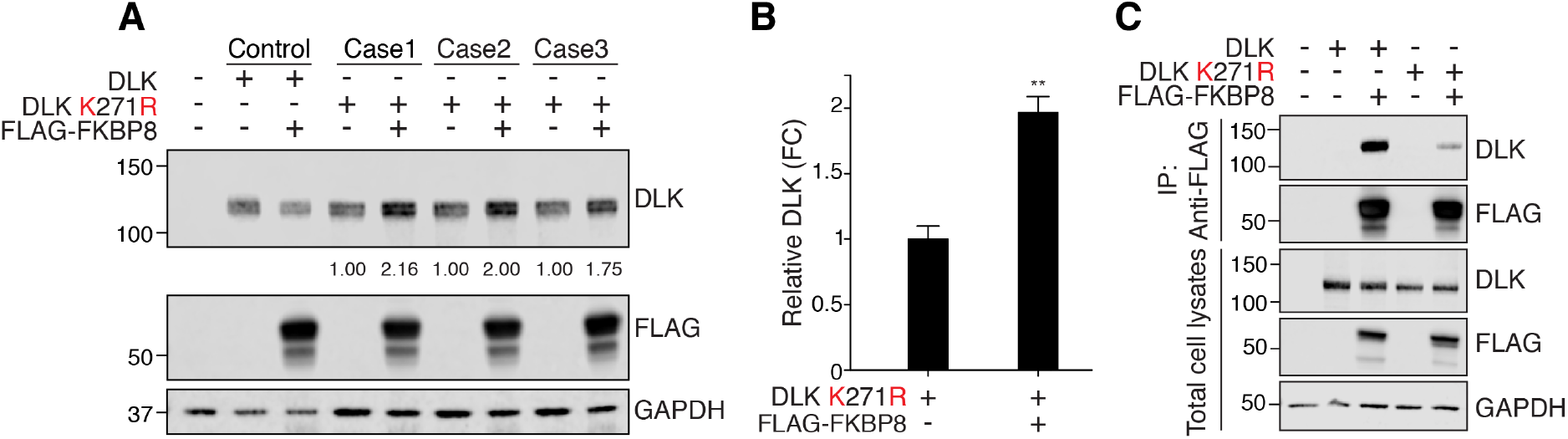
FKBP8 mediates ubiquitin-dependent DLK degradation. (A) Western blot analysis for the expression of a K271R mutant with FKBP8. The numbers indicate the normalized relative intensity. (B) Statistical analysis of (A) (n=3 for each condition; **p<0.01 according to a t-test; mean ± S.E.M.). (C) Western blot analysis for immunoprecipitation assays of FKBP8 with DLK and a K271R mutant.

### In vivo gene delivery of FKBP8 delayed axon degeneration in mouse sciatic nerve and enhanced the viability of RGC neurons after nerve injury

Because FKBP8 was the interactor of DLK to regulate DLK degradation, we monitored neuronal injury responses with overexpressing FKBP8 protein in vivo because DLK is a core regulator of signal transductions for axon regeneration and degeneration. In vivo gene delivery using an adeno-associated virus (AAV) successfully expressed FKBP8 protein in DRG tissues (Figures 6A, 6B and 6C). To test axon degeneration in vivo, the mouse sciatic nerve was cut, and the distal part was dissected at 3 days after axotomy (Figure 6A). Immunohistological analysis showed that FKBP8 overexpression delayed axon degeneration in sciatic nerves because cross-sectioned sciatic nerves of the distal to the cut sites had more TUJ1-positive axons in FKBP8-overexpressing mice three days after axotomy (Figures 6D and 6E). By assessing the number of axonal cross section with TUJ1-positive immunostaining, control sciatic nerves had an average of 11.8 ± 2.4 intact axons with a diameter of more than 5 μm per unit area, while sciatic nerves from FKBP8-overexpressing mice had an average of 20.0 ± 3.2 axons per unit area, a nearly two-fold increase. However, FKBP8-overexpression did not change the efficiency of axon regeneration in the sciatic nerve. Axon regeneration was assessed by SCG10 immunostaining the longitudinal sections of sciatic nerves crushed and dissected at 3 days after injury. SCG10-positive regenerating axons in the sciatic nerves showed no significant difference between control and FKBP8-overexpressing mice (Figures 6F and 6G). These results implied that FKBP8-mediated injury responses might be related to degeneration processes more effectively than regeneration.

**Figure 6.**
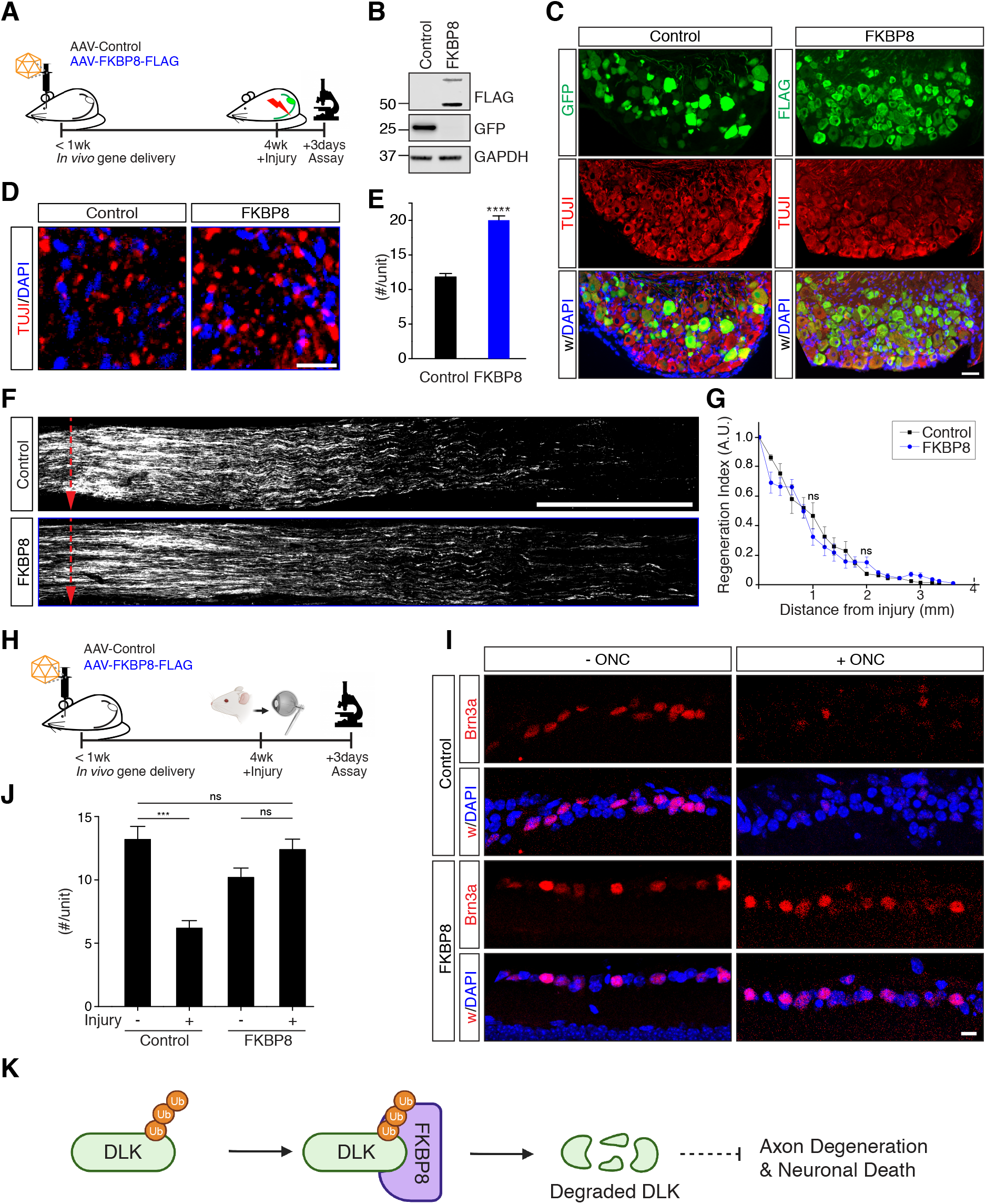
In vivo gene delivery of FKBP8-delayed axon degeneration and enhanced viability of RGC neurons. (A) Experimental scheme for in vivo gene delivery used in in vivo axon regeneration and degeneration assays for mouse sciatic nerves (wk, week). (B) Western blot analysis of GFP or FKBP8 protein from DRG tissue dissected from AAV-injected mice. (C) Immunohistochemistry of mouse DRG sections from AAV-injected mice, stained with anti-GFP for GFP injected mice and anti-FLAG for FKBP8 injected mice. Scale bar, 50μm. (D) In vivo degeneration assays for sciatic nerves. Representative cross-sections of the sciatic nerves from control or FKBP8-expressing mice. Scale bar, 25 μm. (E) Statistical analysis for (D) (n=3 for each condition; ****p≤0.0001 from a *t*-test; mean ± S.E.M.). (F) In vivo axon regeneration assays for sciatic nerves. Representative longitudinal sections of the sciatic nerves from control or FKBP8-expressing mice. The red dotted arrows indicate the injury site. Scale bar, 1 mm. (G) In vivo regeneration index from (F) (n=3 for the control, 6 for FKBP8; ns, not significant from a *t*-test; mean ± S.E.M.). (H) Experimental scheme for in vivo gene delivery used in in vivo axon degeneration assays for mouse retinas (wk; week). (I) Representative longitudinal sections of the retinas from control or FKBP8-expressing mice. Scale bar, 10 μm. (J) Quantification of the number of Brn3a-stained RGCs with or without injury (n=5 for each condition; ***p<0.001; ns, not significant from a *t*-test; mean ± S.E.M.). (K) Schematic illustration of FKBP8-mediated DLK degradation.

Because DLK is responsible for retinal ganglion cell (RGC) apoptosis after optic nerve crush (ONC) injury (Larhammar *et al*, 2017; Huntwork-Rodriguez *et al*, 2013), we tested if FKBP8 overexpression protected against RGC death. Longitudinal sections of mouse retinas were prepared three days after the ONC injury and immunostained with Brn3a as a marker of RGCs (Figure 6H). We observed a significant reduction in Brn3a-positive RGCs in the retina with an ONC injury compared to control mice (Figure 6I). Control mice had an average of 13.2 ± 1.0 Brn3a-positive cells, while the ONC injury reduced this to 6.2 ± 0.6 per unit area. However, the FKBP8-overexpressing mice had an average of 12.4 ± 0.8 at three days after the optic nerve injury (Figure 6J). These results showed that injury-induced retrograde signaling responsible for RGC death after optic nerve crush was downregulated by FKBP8 overexpression. Altogether, FKBP8 might be a potential target for understanding injury-related axon degeneration and neuronal death.

## Discussion

DLK is a core protein responsible for injury-responses and directs neuronal fates under various types of stresse conditions. Because DLK plays a role in both axon regeneration and degeneration, DLK has been referred to as a “double-edged sword” in the reconstruction of damaged neural tissue (Tedeschi & Bradke, 2013). Therefore, it is essential to determine the molecular mechanisms regulating DLK functions for understanding neuronal responses to stresses. In this study, we presented the DLK-interacting proteins FKBPL and FKBP8 as regulators of DLK degradation and DLK kinase activity. FKBPL and FKBP8 bound to the kinase domain of DLK and inhibited DLK kinase activity. In addition, FKBP8 induced the degradation of ubiquitinated DLK through lysosomal degradation pathways. In vivo gene delivery of FKBP8 delayed axon degeneration in sciatic nerves after axotomy and showed a protective effect against RGC death after an optic nerve crush injury, consistent with the suppression of DLK function that promotes axon degeneration and injury-induced neuronal death (references).

DLK protein levels are differentially regulated when neurons are subject to specific stimulations such as axotomy and microtubule stabilizing or destabilizing agents (Fernandes *et al*, 2014; Summers *et al*, 2020; Valakh *et al*, 2015; Jin & Zheng, 2019; Geden & Deshmukh, 2016; DiAntonio, 2019). Moreover, elevated levels of DLK proteins result in neuronal death in optic nerve injury models (Larhammar *et al*, 2017; Huntwork-Rodriguez *et al*, 2013). Therefore, detailing the molecular mechanisms for DLK protein turnover is important for understanding how the fates of neurons are determined in response to injury. Phr1 E3 ligase and the de-ubiquitinating enzyme USP9X are key players regulating DLK protein levels (Babetto *et al*, 2013). Here, we extend the knowledge about DLK protein degradation by identifying the specific lysine residue, lysine 271, responsible for ubiquitination in the kinase domain. Lysine 271 is also responsible for DLK sumoylation, indicating that this residue may be the site of competition between ubiquitination and sumoylation, which is a new finding in terms of DLK post-translational modifications and the mechanisms potentially regulating DLK functions. The lysine 271 site is required for ubiquitin-dependent DLK degradation via lysosomal functions. Therefore, the FKBPL- and FKBP8-mediated DLK protein degradation pathway with lysosomal autophagic functions provides a new direction for the manipulation of DLK protein levels in vivo under neuropathological and neurodegenerative conditions.

## Materials and Methods

### Mice and surgical procedures

CD-1: Crl:CD1(ICR) and C57BL/6J mice were used in the present study. All animal husbandry and surgical procedures were approved by the Korea University Institutional Animal Care & Use Committee (AU-IACUC). Surgery was performed under isoflurane anesthesia following regulatory protocols. Sciatic nerve injury experiments were performed as previously described (Cho *et al*, 2014). Briefly, anesthetized animals were subjected to unilateral exposure of the sciatic nerve at thigh level and a crush injury was inflicted with fine forceps for 10 s.

### Lentiviral constructs and AAV-mediated in vivo gene delivery

Lentivirus-mediated gene delivery was used to knock down target mRNA from embryonic DRG neurons. Lentivirus was produced with Lenti-X packaging Single Shots (Takara, 631275) as previously described (Cho & Cavalli, 2012). For in vitro gene delivery, lentivirus was applied to embryonic DRG neuron cultures at DIV2. To knock down DLK in vitro, shRNA targeting sequences identified by the BROAD Institute (TRCN0000322150) were synthesized (Bionics) and ligated into a pLKO.1 lentiviral vector with the restriction sites AgeI/EcoRI. Lentivirus was produced using Lenti-XTM Packaging Single Shot (Qiagen, 631276), concentrated using a Lenti-XTM Concentrator (Qiagen, 631232), and quantified using a Lenti-XTM GoStixTM Plus kit (Qiagen, 631280) as previously described (Jeon *et al*, 2021). The efficiency of the knockdown process was confirmed using RT-qPCR. To deliver genes in vivo, 10 μl of adeno-associated virus (AAV, serotype 9)-encoding mouse Flag-tagged FKBP8 was injected into neonatal CD-1 mice (postnatal day 1) via a facial vein injection using a Hamilton syringe (Hamilton, 1710 syringe with a 33G/0.75-inch small hub removable needle). The expression of GFP and the target genes in the sciatic nerve and DRGs was confirmed with immunoblot, immunohistochemistry, and RT-qPCR analysis.

### Yeast two-hybrid screening

Yeast two-hybrid (Y2H) analysis was performed using a contract with Panbionet (http://panbionet.com/) as described previously. The bait was generated from mouse Map3k12 CDS (DLK, NM_001163643, full length, 887 amino acids, 2,667) and cloned into Xmal/SalI sites of a pGBKT7 vector with the primers 5’-CGC CCG GGG GCC TGC CTC CAT GAA ACC C-3’ and 5’- GG CTC GAG TCA TGG AGG AAG GGA GGC T-3’. Various DLK baits were used to screen multiple cDNA libraries derived from mouse embryos.

### Antibodies and chemicals

The following antibodies were used: anti-GFP (Santa Cruz, sc-9996 for co-immunoprecipitation; Abcam, ab32146 for immunoblots), anti-Flag HRP-conjugated (Sigma, A8592), anti-p-SEK1/MKK4 (Cell Signaling, CST-9151), anti-GST (Santa Cruz, sc-138), anti-alpha tubulin (Santa Cruz, sc-53030), anti-DLK (ThermoFisher Scientific, PA5-32173 for co-immunoprecipitation; Antibodies Incorporated, 75-355 for immunoblot), anti-p-SAPK/JNK (Cell Signaling, CST-9251S), anti-GAPDH (Santa Cruz, sc-32233), anti-p-cJun (Cell Signaling, CST-9251S), anti-LC3A/B (Cell Signaling, CST-12741), anti-SCG10 (Novus Biologicals, NBP1-49461), and anti-beta III tubulin (Abcam, ab41489). We dissolved all chemicals in DMSO (Sigma, D8418-250ML) and treated the controls with this vehicle except vincristine (Sigma, V8879), which was dissolved in methanol. We used vincristine at 200 nM, bafilomycin (Sigma, B1793) at 100 nM, caspase inhibitor (Sigma, 400012) at 1 and 5 μM, pan-caspase inhibitor (R&D systems, FMK001) at 10 and 50 μM, and MG-132 (Sigma, M7449) at 10 μM.

### In vitro degeneration assays

Embryonic DRGs were cultured in 12-well plates (SPL) coated with poly-d-lysine/laminin. For lentiviral transduction, lentivirus was added to the culture at DIV2. The culture medium was changed at DIV5, then vincristine and bafilomycin were added at DIV7. Within 48 hours, images were taken every 12 hours for the degeneration assays. Axon degeneration was analyzed as described previously. Briefly, phase-contrast images were obtained using an inverted light microscope (CKX53; Olympus). Three non-overlapping images of each well were taken at each time point and were assessed for axon degeneration. Images were processed with the auto-level function in Photoshop (Adobe) for brightness adjustment. The images were then analyzed using a macro written in ImageJ to calculate the degeneration index (Araki *et al*, 2004; Shin *et al*, 2012; Miller *et al*, 2009). After the images were binarized, the total axon area was defined by the total number of detected pixels. The area of degenerated axon fragments was calculated using the particle analyzer function. To calculate the degeneration index, we divided the area covered by the axon fragments by the total axon area. The average of three images taken from the same well was used to calculate the mean degeneration index for each well.

### Optic nerve injury and retina tissue preparation

To expose the optic nerve, the conjunctiva from the orbital region of the eye was cleared then the optic nerve was crushed for 3 seconds with Dumont #5 forceps (Fine Science Tools, 11254-20) with special care taken not to damage the vein sinus. A saline solution was applied before and after the optic nerve crush injury to protect the eye from desiccation. Three days after injury, the mouse eyes were dissected and fixed via immersion in a 4% paraformaldehyde solution for 2 hours. After being washed three times in PBS, the eyes were transferred to 30% sucrose solution for 24 hours at 4°C. The optic nerves were then dissected out with micro-scissors (Fine Science Tools, 15070-08), sectioned at 15 μm in a cryostat, immunostained with Brn3a, and mounted in the mounting medium VectaShield (Vector Laboratories, H1000 or H1200).

### Immunocytochemistry and immunohistochemistry

Cultured neurons were fixed in 4% paraformaldehyde for 20 minutes at room temperature. Samples were washed with 0.1% Triton X100 in PBS (PBS-T) and immunostained using the same procedure described for the immunohistochemistry. To measure the axonal length, samples stained with anti-beta III tubulin antibody were imaged with an EVOS FL Auto 2 microscope and a Zeiss LSM 800 confocal microscope. DRG and sciatic nerve tissues were fixed immediately after dissection in 4% paraformaldehyde for 1 hour at room temperature and immersed in 30% sucrose. Samples were cryopreserved in OCT medium (Tissue-Tek), cryo-sectioned at a thickness of 10 μm, and immunostained as described previously. Briefly, samples were blocked in blocking solution (5% normal goat serum and 0.1% Triton X-100 in PBS) for 1 hour and incubated with primary antibodies diluted in blocking solution overnight at 4°C. Samples were then rinsed twice with 0.1% Triton X100 in PBS (PBS-T), incubated with secondary antibodies for 1 hour at room temperature, rinsed three times with PBS-T, and mounted in VectaShield (Vector Laboratories, H1000 or H1200). The samples were imaged with an EVOS FL Auto 2 microscope (Thermo, AMAFD2000) or a Zeiss LSM800.

### In vivo axon regeneration assays

To examine axon regeneration in sciatic nerves, sciatic nerves were dissected three days after the crush injury and dissected, sectioned, and immunostained with TUJ1 and anti-SCG10 antibodies. Immunostained sections were imaged with an EVOS FL Auto 2 Imaging System (Thermo, AMAFD2000), which automatically combined the individual images. The fluorescence intensity of SCG10 was measured along the length of the nerve section using ImageJ software. A regeneration index was calculated by measuring the average SCG10 intensity from the injury site to the distal side normalized to the intensity at the crush site and presented as a regeneration index.

### In vivo degeneration assays

For the axon degeneration assays, the sciatic nerves of FKBP8 overexpressed mice were dissected three days after the crush injury and dissected and immunostained with TUJ1 antibody. Immunostained sections were imaged with an EVOS FL Auto 2 Imaging System and a Zeiss LSM800. We quantified the unfragmented axons in the distal nerve and compared these between FKBP8-overexpressed nerves and the control.

### Western blot analysis and co-immunoprecipitation assays

To study protein–protein interactions, plasmids containing mouse DLK and mouse Flag-tagged FKBPL/FKBP8 were transfected into HEK293T cells using Lipofectamine 2000 (Thermo, 11668-019) following the manufacturer’s instructions. Cell lysates were prepared in 1X SDS buffer (63 mM Tris pH 6.8, 2% SDS, 10% glycerol) then boiled for 10 minutes at 95°C. After centrifugation, the protein concentration in the supernatant was determined using DC protein assays (Bio-rad, 5000116) with bovine serum albumin solutions as standards. Equal amounts of protein were loaded into 1x MOPS running buffer for SDS-PAGE and transferred to a nitrocellulose membrane. The membranes were blocked with 5% skim milk dissolved in 1x TBS with 0.1% Tween-20 (TBS-T) for 1 hour, incubated with primary antibodies overnight at 4°C, and washed three times with TBS-T. The blots were then incubated with secondary antibodies for 1 hour and washed three times with TBS-T. Protein expression levels were analyzed with enhanced chemiluminescence using Odyssey (Li-Cor).

DLK and Flag-tagged FKBPL/FKBP8 transfected HEK293T cells were lysated in IP buffer (0.5% NP40, 150 mM NaCl, 20 mM Tris-HCl pH 7.5) containing a protease inhibitor cocktail (Roche). GFP-DLK was immunoprecipitated with anti-GFP antibody pre-bound to Dynabeads Protein A (Thermo, 10001D) from input lysates for 16 hours at 4°C. The precipitants were washed four times using DynaMag-2 (Thermo, 12321D) and subjected to SDS-PAGE for western blot analysis.

### In vitro kinase assays

DLK kinase activity was assessed as described previously (Holland *et al*, 2016). HEK293T cells were transfected with GFP-tagged DLK or FLAG-tagged FKBPL individually using Lipofectamine 2000 (Thermo, 11668-019). Cell lysates were prepared in immunoprecipitation buffer (50 mM HEPES, pH 7.5, 150 mM NaCl, 1 mM EGTA, 0.1% Triton X-100) containing a protease inhibitor cocktail (Roche, 11836153001). GFP-DLK was immunopurified using anti-GFP antibody with Dynabeads Protein G (ThermoFisher Scientific, 10007D). The substrate GST-MKK4 was purified following a previous protocol (Holland *et al*, 2016). Complexes were incubated for 30 min at 30 °C in 30 μl of kinase buffer (25 mM HEPES, pH 7.2, 10% glycerol, 100 mM NaCl, 20 mM MgCl2, 0.1 mM sodium vanadate, and protease inhibitors) containing 25 μM ATP and 2 μg of GST or GST-MKK4. Reactions were terminated by the addition of Laemmli buffer, boiled, resolved using SDS-PAGE, and subjected to western blot analysis with anti-phospho-MKK4 antibody.

## Acknowledgements

This work was supported by a National Research Foundation of Korea (NRF) grant funded by the Korean government (MSIT) (NRF-2019R1A2C1005380 to Y.C, 2016R1A5A2007009 and 2020R1C1C1011074 to J.E.S.) and by a Korea University grant.

## Author contributions

Conceptualization, B.L., J.E.S., A.D., V.C. and Y.C.; Methodology, Formal Analysis, and Investigation, B.L., Y.O., E.C., J.E.S. and Y.C.; Resources, Funding Acquisition, Y.C. and J.E.S; Supervision and Project Administration, J.E.S., A.D., V.C. and Y.C.; Data Curation and Visualization, B.L. and Y.C.; Writing – Original Draft, B.L. and Y.C.; Writing – Review & Editing, B.L., J.E.S., A.D., V.C., and Y.C.

## Data Availability Section

All data is available in the main text.

## References

Araki T, Sasaki Y & Milbrandt J (2004) Increased nuclear NAD biosynthesis and SIRT1 activation prevent axonal degeneration. Science 305: 1010–1013

Asghari Adib E, Smithson LJ & Collins CA (2018) An axonal stress response pathway: degenerative and regenerative signaling by DLK. Curr Opin Neurobiol 53: 110–119

Babetto E, Beirowski B, Russler E V., Milbrandt J & DiAntonio A (2013) The Phr1 Ubiquitin Ligase Promotes Injury-Induced Axon Self-Destruction. Cell Rep 3: 1422–1429

Bhujabal Z, Birgisdottir ÅB, Sjøttem E, Brenne HB, Øvervatn A, Habisov S, Kirkin V, Lamark T & Johansen T (2017) FKBP8 recruits LC3A to mediate Parkin-independent mitophagy. EMBO Rep 18: 947–961

Chen X, Rzhetskaya M, Kareva T, Bland R, During MJ, Tank AW, Kholodilov N & Burke RE (2008) Antiapoptotic and trophic effects of dominant-negative forms of dual leucine zipper kinase in dopamine neurons of the substantia nigra in vivo. J Neurosci 28: 672–680

Cho Y & Cavalli V (2012) HDAC5 is a novel injury-regulated tubulin deacetylase controlling axon regeneration. EMBO J 31: 3063–3078

Cho Y, Di Liberto V, Carlin D, Abe N, Li KH, Burlingame AL, Guan S, Michaelevski I & Cavalli V (2014) Syntaxin13 expression is regulated by mammalian target of rapamycin (mTOR) in injured neurons to promote axon regeneration. J Biol Chem 289: 15820–15832

Cho Y, Sloutsky R, Naegle KM & Cavalli V (2013) Injury-Induced HDAC5 nuclear export is essential for axon regeneration. Cell 155: 894–908

Collins CA, Wairkar YP, Johnson SL & DiAntonio A (2006) Highwire restrains synaptic growth by attenuating a MAP kinase signal. Neuron 51: 57–69

DiAntonio A (2019) Axon degeneration: mechanistic insights lead to therapeutic opportunities for the prevention and treatment of peripheral neuropathy. Pain 160: S17–S22

Fan G, Merritt SE, Kortenjann M, Shaw PE & Holzman LB (1996) Dual leucine zipper-bearing kinase (DLK) activates p46(SAPK) and p38(mapk) but not ERK2. J Biol Chem 271: 24788–24793

Fernandes KA, Harder JM, John SW, Shrager P & Libby RT (2014) DLK-dependent signaling is important for somal but not axonal degeneration of retinal ganglion cells following axonal injury. Neurobiol Dis 69: 108–116

Frey E, Valakh V, Karney-Grobe S, Shi Y, Milbrandt J & DiAntonio A (2015) An in vitro assay to study induction of the regenerative state in sensory neurons. Exp Neurol 263: 350–363

Geden MJ & Deshmukh M (2016) Axon degeneration: Context defines distinct pathways. Curr Opin Neurobiol 39: 108–115

Geisler S, Doan RA, Strickland A, Huang X, Milbrandt J & DiAntonio A (2016) Prevention of vincristine-induced peripheral neuropathy by genetic deletion of SARM1 in mice. Brain 139: 3092–3108

Ghartey-Kwansah G, Li Z, Feng R, Wang L, Zhou X, Chen FZ, Xu MM, Jones O, Mu Y, Chen S, et al (2018) Comparative analysis of FKBP family protein: evaluation, structure, and function in mammals and Drosophila melanogaster. BMC Dev Biol 18: 7

Ghosh AS, Wang B, Pozniak CD, Chen M, Watts RJ & Lewcock JW (2011) DLK induces developmental neuronal degeneration via selective regulation of proapoptotic JNK activity. J Cell Biol 194: 751–764

Heitman J, Movva NR & Hall MN (1992) Proline isomerases at the crossroads of protein folding, signal transduction, and immunosuppression. New Biol 4: 448–460

Hirai SI, De FC, Miyata T, Ogawa M, Kiyonari H, Suda Y, Aizawa S, Banba Y & Ohno S (2006) The c-Jun N-terminal kinase activator dual leucine zipper kinase regulates axon growth and neuronal migration in the developing cerebral cortex. J Neurosci 26: 11992–12002

Holland SM, Collura KM, Ketschek A, Noma K, Ferguson TA, Jin Y, Gallo G & Thomas GM (2016a) Palmitoylation controls DLK localization, interactions and activity to ensure effective axonal injury signaling. Proc Natl Acad Sci U S A 113: 763–768

Holland SM, Collura KM, Ketschek A, Noma K, Ferguson TA, Jin Y, Gallo G & Thomas GM (2016b) Palmitoylation controls DLK localization, interactions and activity to ensure effective axonal injury signaling. Proc Natl Acad Sci 113: 763 LP – 768

Huntwork-Rodriguez S, Wang B, Watkins T, Ghosh AS, Pozniak CD, Bustos D, Newton K, Kirkpatrick DS & Lewcock JW (2013) JNK-mediated phosphorylation of DLK suppresses its ubiquitination to promote neuronal apoptosis. J Cell Biol 202: 747–763

Jeon Y, Shin JE, Kwon M, Cho E, Cavalli V & Cho Y (2020) In Vivo Gene Delivery of STC2 Promotes Axon Regeneration in Sciatic Nerves. Mol Neurobiol

Jin Y & Zheng B (2019) Multitasking: Dual leucine zipper-bearing kinases in neuronal development and stress management. Annu Rev Cell Dev Biol 35: 501–521

Kang CB, Hong Y, Dhe-Paganon S & Yoon HS (2008) FKBP family proteins: immunophilins with versatile biological functions. Neurosignals 16: 318–325

Larhammar M, Huntwork-Rodriguez S, Jiang Z, Solanoy H, Ghosh AS, Wang B, Kaminker JS, Huang K, Eastham-Anderson J, Siu M, et al (2017) Dual leucine zipper kinase-dependent PERK activation contributes to neuronal degeneration following insult. Elife 6: 1–27

Lee B, Lee J, Jeon Y, Kim H, Kwon M, Shin JE & Cho Y (2021) Promoting axon regeneration by enhancing the non-coding function of the injury-responsive coding gene *Gpr151*. bioRxiv: 2021.02.19.431965

Lim GG & Lim K-L (2017) Parkin-independent mitophagy-FKBP8 takes the stage. EMBO Rep 18: 864–865

Martin DDO, Kanuparthi PS, Holland SM, Sanders SS, Jeong HK, Einarson MB, Jacobson MA & Thomas GM (2019) Identification of Novel Inhibitors of DLK Palmitoylation and Signaling by High Content Screening. Sci Rep 9: 1–12

Miller BR, Press C, Daniels RW, Sasaki Y, Milbrandt J & Diantonio A (2009) A dual leucine kinase-dependent axon self-destruction program promotes Wallerian degeneration. Nat Neurosci 12: 387–389

Misaka T, Murakawa T, Nishida K, Omori Y, Taneike M, Omiya S, Molenaar C, Uno Y, Yamaguchi O, Takeda J, et al (2018) FKBP8 protects the heart from hemodynamic stress by preventing the accumulation of misfolded proteins and endoplasmic reticulum-associated apoptosis in mice. J Mol Cell Cardiol 114: 93–104

Montersino A & Thomas GM (2015) Slippery signaling: Palmitoylation-dependent control of neuronal kinase localization and activity. Mol Membr Biol 32: 179–188

Nakata K, Abrams B, Grill B, Goncharov A, Huang X, Chisholm AD & Jin Y (2005) Regulation of a DLK-1 and p38 MAP kinase pathway by the ubiquitin ligase RPM-1 is required for presynaptic development. Cell 120: 407–420

Niu J, Sanders SS, Jeong HK, Holland SM, Sun Y, Collura KM, Hernandez LM, Huang H, Hayden MR, Smith GM, et al (2020) Coupled Control of Distal Axon Integrity and Somal Responses to Axonal Damage by the Palmitoyl Acyltransferase ZDHHC17. Cell Rep 33: 108365

Ren J, Gao X, Jin C, Zhu M, Wang X, Shaw A, Wen L, Yao X & Xue Y (2009) Systematic study of protein sumoylation: Development of a site-specific predictor of SUMOsp 2.0. Proteomics 9: 3409–3412

Schmid FX, Mayr LM, Mücke M & Schönbrunner ER (1993) Prolyl isomerases: role in protein folding. Adv Protein Chem 44: 25–66

Shin JE, Cho Y, Beirowski B, Milbrandt J, Cavalli V & DiAntonio A (2012a) Dual Leucine Zipper Kinase Is Required for Retrograde Injury Signaling and Axonal Regeneration. Neuron 74: 1015–1022

Shin JE, Ha H, Cho EH, Kim YK & Cho Y (2018) Comparative analysis of the transcriptome of injured nerve segments reveals spatiotemporal responses to neural damage in mice. J Comp Neurol

Shin JE, Ha H, Kim YK, Cho Y & DiAntonio A (2019) DLK regulates a distinctive transcriptional regeneration program after peripheral nerve injury. Neurobiol Dis 127: 178–192

Shin JE, Miller BR, Babetto E, Cho Y, Sasaki Y, Qayum S, Russler E V, Cavalli V, Milbrandt J & DiAntonio A (2012b) SCG10 is a JNK target in the axonal degeneration pathway. Proc Natl Acad Sci U S A 109: E3696–705

Summers DW, Frey E, Walker LJ, Milbrandt J & DiAntonio A (2020) DLK Activation Synergizes with Mitochondrial Dysfunction to Downregulate Axon Survival Factors and Promote SARM1-Dependent Axon Degeneration. Mol Neurobiol 57: 1146–1158

Tedeschi A & Bradke F (2013a) The DLK signalling pathway - A double-edged sword in neural development and regeneration. EMBO Rep 14: 605–614

Tedeschi A & Bradke F (2013b) The DLK signalling pathway - A double-edged sword in neural development and regeneration. EMBO Rep 14: 605–614

Tong M & Jiang Y (2015) FK506-Binding Proteins and Their Diverse Functions. Curr Mol Pharmacol 9: 48–65

Valakh V, Frey E, Babetto E, Walker LJ & DiAntonio A (2015a) Cytoskeletal disruption activates the DLK/JNK pathway, which promotes axonal regeneration and mimics a preconditioning injury. Neurobiol Dis 77: 13–25

Valakh V, Frey E, Babetto E, Walker LJ & DiAntonio A (2015b) Cytoskeletal disruption activates the DLK/JNK pathway, which promotes axonal regeneration and mimics a preconditioning injury. Neurobiol Dis 77: 13–25

Watkins TA, Wang B, Huntwork-Rodriguez S, Yang J, Jiang Z, Eastham-Anderson J, Modrusan Z, Kaminker JS, Tessier-Lavigne M & Lewcock JW (2013) DLK initiates a transcriptional program that couples apoptotic and regenerative responses to axonal injury. Proc Natl Acad Sci U S A 110: 4039–4044

Welsbie DS, Schirok H, Mitchell K, Koch M, Kim B-J, Lobell M, Patel AK, Holton S, Hristodorov D, Esteve-Rudd J, et al (2018) Identification of a retinal neuroprotective kinase inhibitor with preferential activity against DLK compared to LZK. Invest Ophthalmol Vis Sci 59: 2493

Yoo S-M, Yamashita S, Kim H, Na D, Lee H, Kim SJ, Cho D-H, Kanki T & Jung Y-K (2020) FKBP8 LIRL-dependent mitochondrial fragmentation facilitates mitophagy under stress conditions. FASEB J 34: 2944–2957

Zhao Q, Xie Y, Zheng Y, Jiang S, Liu W, Mu W, Liu Z, Zhao Y, Xue Y & Ren J (2014) GPS-SUMO: a tool for the prediction of sumoylation sites and SUMO-interaction motifs. Nucleic Acids Res 42: W325–W330

